# Protocol for mapping the spatial variability in cell wall mechanical bending behavior in living leaf pavement cells

**DOI:** 10.1101/2021.02.23.432478

**Authors:** Wenlong Li, Sedighe Keynia, Samuel A. Belteton, Faezeh Afshar-Hatam, Daniel B. Szymanski, Joseph A. Turner

## Abstract

An integrated, experimental-computational approach is presented to analyze the variation of elastic bending behavior in the primary cell wall of living *Arabidopsis thaliana* pavement cells and to measure turgor pressure in the cells quantitatively under different osmotic conditions. Mechanical properties, size and geometry of cells and internal turgor pressure greatly influence their morphogenesis. Computational models of plant morphogenesis require values for wall elastic modulus and turgor pressure but very few experiments were designed to validate the results using measurements that deform the entire thickness of the cell wall. Because new wall material is deposited from inside the cell, full-thickness deformations are needed to quantify relevant changes associated with cell development. The approach here uses laser scanning confocal microscopy to measure the three-dimensional geometry of a single pavement cell, and indentation experiments equipped with high magnification objective lens to probe the local mechanical responses across the same cell wall. These experimental results are matched iteratively using a finite element model of the experiment to determine the local mechanical properties, turgor pressure, and cell height. The resulting modulus distribution along the periclinal wall is shown to be nonuniform. These results are consistent with the characteristics of plant cell walls which have a heterogeneous organization. This research and the resulting model will provide a reference for future work associated with the heterogeneity and anisotropy of mechanical properties of plant cell walls in order to understand morphogenesis of the primary cell walls during growth and to predict quantitatively the magnitudes/directions of cell wall forces.

**One sentence summary:** The distribution of elastic modulus of the periclinal cell walls of living

*Arabidopsis* epidermis is nonuniform as measured by bending the entire thickness of the wall.

**Highlights:**

- Experimental characterization of the spatial distribution of elastic bending behavior across the periclinal wall
- Quantification of the turgor pressure of the living plant epidermal cells validated with osmotic treatments
- Quantification of the effect of cell geometry on the measured mechanical response

**Graphical abstract:** 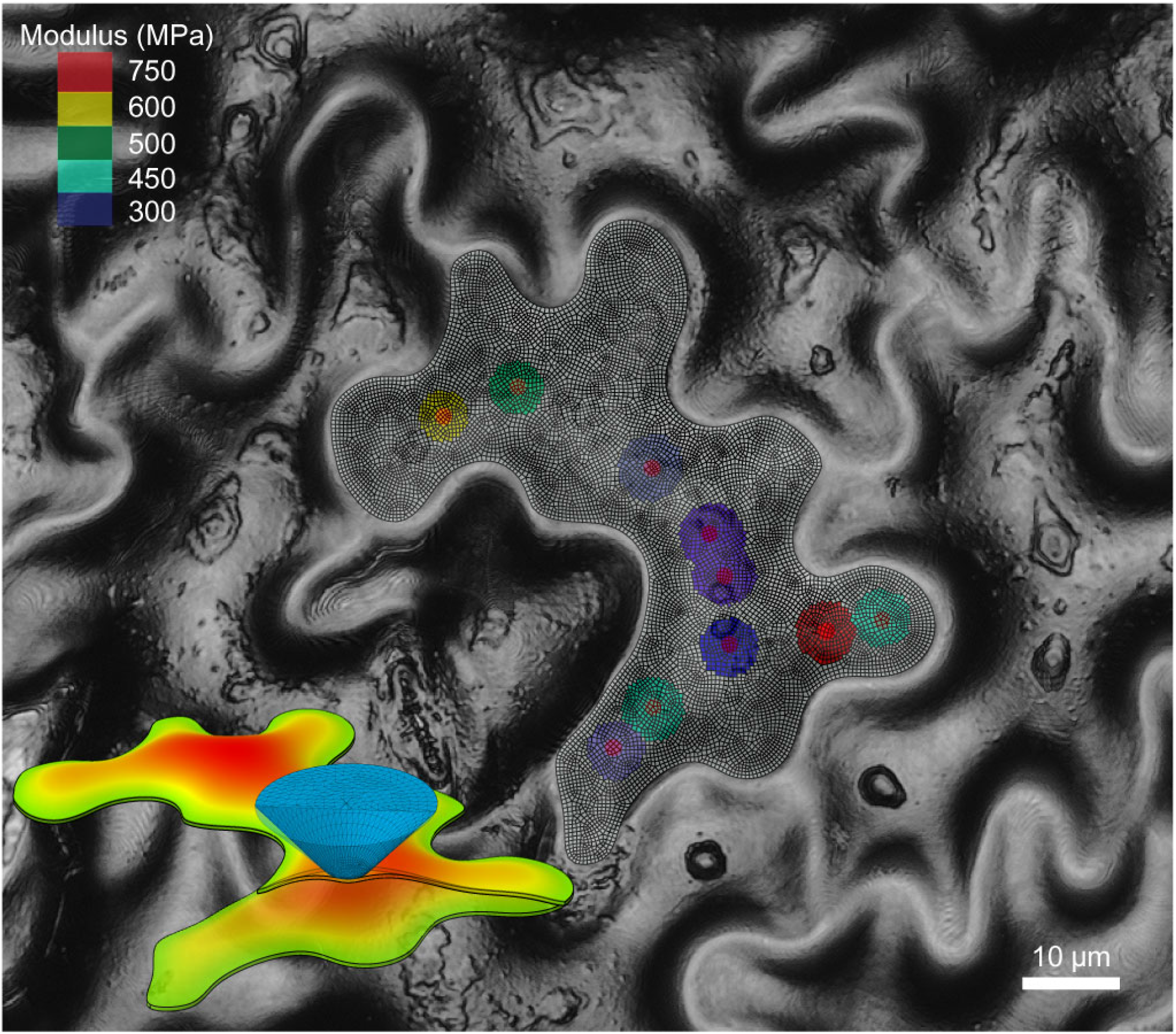

## Introduction and Motivation

The plant leaf epidermis consists of a single layer of cells that adhere to one another through their middle lamella. Many species have jigsaw-puzzle-piece shaped cells and grow with an interdigitated growth pattern (Panteris and Galatis, 2005; Vőfély et al., 2019). This behavior is attributed to the anisotropic enlargement of cells as they expand (Panteris and Galatis, 2005; Szymanski, 2014). The formation of this patterning has attracted much attention in order to discover the fundamental mechanisms that govern the cell shapes. Mechanical properties such as the cell wall elastic modulus and its distribution are clearly important and mechanical structural analysis is used for that reason (Cosgrove, 1993; Sampathkumar et al., 2014; Majda et al., 2017; Sapala et al., 2018; Bidhendi et al., 2019). In some cases, a qualitative analysis is sufficient to uncover trends in cell growth. However, quantitative information about the stresses and strains in the cell walls is critical if growth mechanisms are to be translated temporally and across different species. Computational models are critical to such analyses because they can provide potential mechanisms that can be explored experimentally. Quantitative knowledge of the wall elastic modulus tensor is especially critical because it provides the connection between the wall stresses and strains. Erroneous conclusions about growth mechanisms may result if computational models are not based on accurate material behavior. As a result, it is necessary to develop a reliable approach to measure the cell wall mechanical properties and turgor pressure\ accurately in these living systems. An example of pavement cell lobing is illustrated in Fig. 1A-C in which a single cell is tracked temporally (9.5 hours) in order to quantify the space-time characteristics of the lobing wall indicated by the arrows (see Belteton et al., 2021 for details about imaging and more examples). Although many features for this cell appear self-similar (only larger), the forming lobes become deeper with a spatial variability that is difficult to predict from the initial state. Many efforts have been made to understand the biomechanics behind this type of patterning which is an integral part of morphogenesis(Cosgrove, 1993; Sampathkumar et al., 2014; Majda et al., 2017; Sapala et al., 2018; Bidhendi et al., 2019; Lin and Yang, 2020; Belteton et al., 2021)

**Figure 1.**
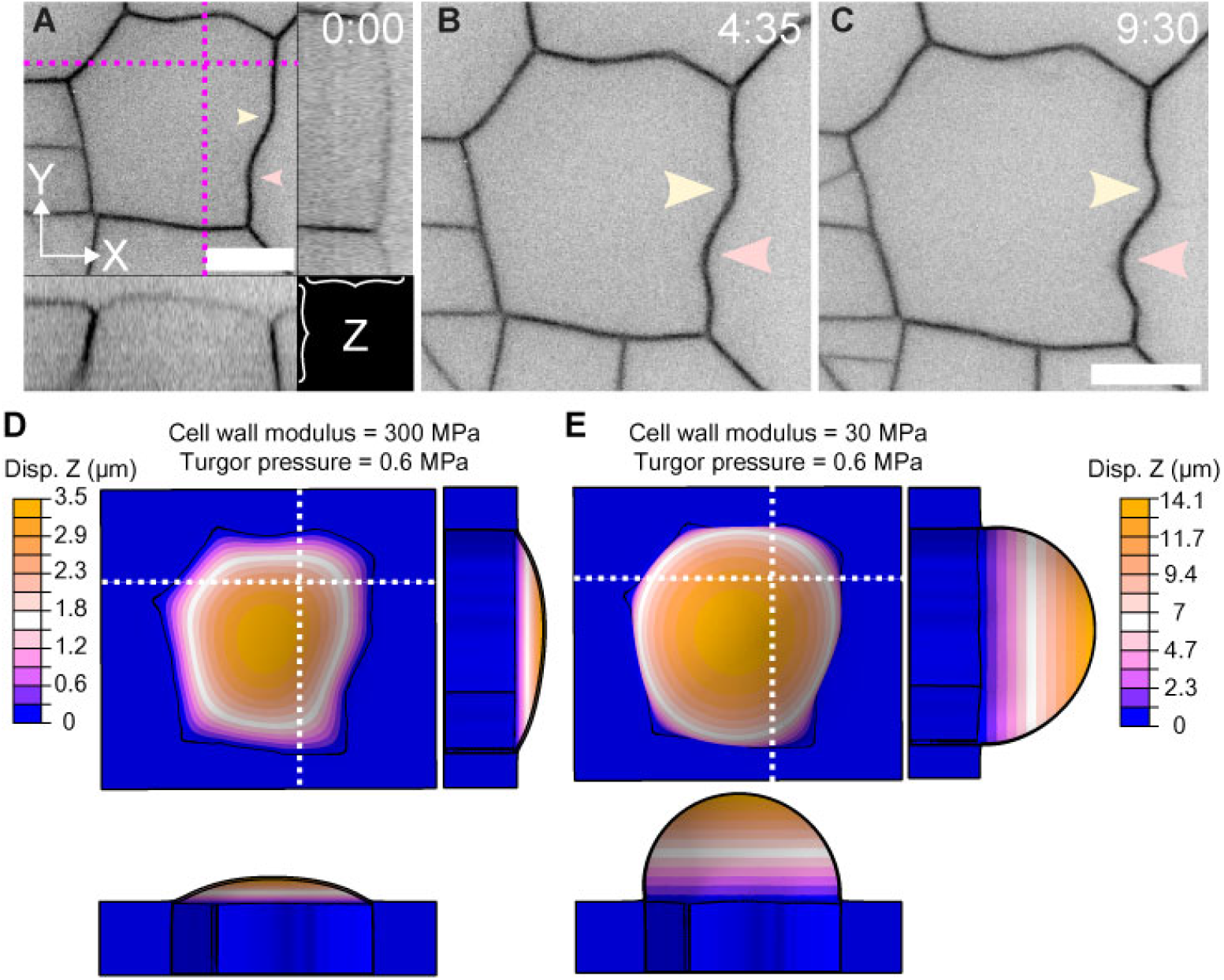
Heterogeneous growth and example FE analysis showing the impact of cell wall modulus. (A-C) Morphogenesis of lobe features detected during time-lapsed imaging, lobe locations are indicated with yellow and pink arrows, and the time intervals in hours are shown. Insets in panel A show the y-z (right) and x-z (below) views of the same cell to reveal the outer periclinal and anticlinal walls. Scale bar for (A-C) = 10 µm. (D-E) FE analysis is conducted for the same cell in A using two values of uniform cell wall modulus (wall thickness = 300 nm; turgor pressure = 0.6 MPa). Cross-sectional views of the y-z and x-z are shown for comparison with the experiments.

Computational models are often used for such studies, but there is little agreement in the literature regarding the appropriate mechanical properties that should be used in models. The impact of the modulus value on the wall deformation behavior is illustrated in Fig. 1D-E. The cell shown in Fig. 1A is modeled using the finite element (FE) method (details later and in Methods) using two different values for the modulus of the periclinal wall with all other properties constant (thickness = 300 nm; turgor pressure = 0.6 MPa). In Fig. 1D, with a modulus of 300 MPa, the periclinal wall has a maximum height of ∼3.5 μm and the maximum is centered within the cell. In this case, the out-of-plane maximum height is similar to the microscopy results. If the value of the modulus is reduced by a factor of ten (30 MPa), as shown in Fig. 1E, the height is considerably larger (∼14.1 μm), and is well above the height observed in typical pavement cells. The cross-sections of the periclinal wall for the cell and model, shown at the positions of the dashed lines, are also revealing. The *x*-*z* microscopy profile of the cell shows a maximum that is not centered between the two anticlinal walls in contrast to the *y*-*z* profile which has a centered maximum. Both profiles of the model have a centered maximum suggesting that the uniform, isotropic properties or boundary conditions used in the model may be too simplistic. This comparison of simulated cell wall deformation under turgor pressure using different orders of magnitude of cell wall modulus motivates the focus of this article: Can the spatial variability of the periclinal wall and turgor pressure be mapped quantitatively using measurements that deform the entire thickness of the wall in conjunction with a computational model created from the full 3D wall geometry? Variations of both modulus and turgor pressure have been discussed previously.

The possibility of a nonuniform distribution of cell wall mechanical properties has been considered by others conceptually (Szymanski and Cosgrove, 2009; Sampathkumar et al., 2014; Majda et al., 2017; Bidhendi et al., 2019; Hamant et al., 2019). However, a strong connection between anisotropic growth and such a spatial variability is rarely discussed quantitatively due to a lack of experimental data that engages the full thickness of the cell wall. As discussed previously (Cosgrove, 2016), AFM measurements on the outer surface of cell walls often use forces that compress only the outer portion of the wall. Thus, such measurements may not be sensitive to material behavior closer to the interior wall surface. As cells grow, new wall material is added from the inside of the cell. Therefore, changes to wall properties are expected from the inside and measurements that are sensitive to this portion of the wall are needed to capture changes during growth. The ability to quantify and to map the mechanical properties spatially across individual cells can provide needed information for models that are used to establish the relations between stress patterns and growth (e.g., Belteton, et al., 2021).

There have been many attempts to characterize the mechanical properties of plant epidermal cell walls directly, including tensile tests of artificial composites (Chanliaud et al., 2002), micro-testing of peeled/cut leaf samples (Hiller et al., 1996; Sanati Nezhad et al., 2013; Zamil et al., 2015), atomic force microscope (AFM) measurements (Peaucelle, 2014; Sampathkumar et al., 2014; Beauzamy et al., 2015; Carter et al., 2017; Long et al., 2020), and micro/nanoindentation measurements (Hayot et al., 2012; Forouzesh et al., 2013; Weber et al., 2015; Malgat et al., 2016). These different approaches have resulted in a very wide range of cell wall elastic moduli, from 1 MPa (Majda et al., 2017), ∼10 MPa (Sampathkumar et al., 2014), to hundreds of MPa (Forouzesh et al., 2013; Malgat et al., 2016). Some differences in properties might be expected due to the influence of plant age, cell type, and environmental factors. In addition, the organization of the wall is such that the length scale and time scale of each measurement plays a role in the outcome (Cosgrove, 2016). For example, an AFM tip with radius on the order of tens of nanometers will engage a volume of material less than ∼ 1 μm^3^ while micro/nanoindentation measurements use larger tips such that the contact radius is a few micrometers. Instrumented indentation testing (IIT), also called nanoindentation (Fischer-Cripps, 2011), unlike AFM, can engage bending of the full thickness of the cell wall. Variations within a single cell, as determined from measurements of wall bending, would indicate possible organizational heterogeneity in the cell wall constituents – a substantive advance from current knowledge.

Turgor pressure is often thought to be uniform in all cells in a given leaf but few studies have investigated spatial variations quantitatively. Recently, (Long et al., 2020) suggested that turgor pressure could vary within the cells of the shoot apical meristem of *Arabidopsis*. Thus, it also is plausible that differential pressure may exist among pavement cells that can influence the formation of puzzle-shaped cells when combined with the effect of nonuniform mechanical properties. The presence of turgor pressure is a challenge for characterization of mechanical properties of the cell wall due to the coupling effect of the mechanical response when the cell walls are loaded with sufficient loads. Plasmolysis is the most accepted method for measuring turgor pressure. However, many other methods have been proposed including micro/nanoindentation (Forouzesh et al., 2013; Weber et al., 2015; Malgat et al., 2016), probe contact (Lintilhac et al., 2000), pressure chamber (Tyree and Hammel, 1972) and the pressure probe (Wang et al., 2006). Unfortunately, the sizes of cells and cell geometry/topography can limit the viability of some methods for all plant species, especially if a nondestructive approach is needed.

In this article, an iterative experimental-computational approach is used in which IIT and laser confocal scanning microscope (LSCM) measurements are combined with the finite element (FE) method, to quantify the local mechanical properties of periclinal cell walls and turgor pressure of *Arabidopsis thaliana*. High-resolution imaging is used to position the IIT experiments, such that several measurements can be made over the surface of the periclinal wall of the same cell. The three dimensional (3D) topography of the same cell is also mapped with LSCM and used to interpret the IIT data. Our previous IIT research on plant cells (Forouzesh et al., 2013) did not use high resolution imaging, nor was the precise cell geometry (cell shape, cell height) used for analysis of the experimental data. With these mechanical and geometric measurements, the elastic modulus and the associated turgor pressure at each indentation position along the periclinal wall are estimated using the computational model in an iterative manner. Turgor pressure estimates are validated using the same approach for a cell under different osmotic conditions to observe the associated changes. This article describes an important breakthrough technology to map the subcellular distribution of mechanical properties of the periclinal wall nondestructively using deformations that engage the entire thickness of the cell wall. This information is necessary to generate plausible FE models that can simulate stress distributions in the cell wall quantitatively in order to understand the biomechanics of morphogenesis.

## Results

The approach used two types of measurements. A laser scanning confocal microscope (LSCM) (0.5 nm z-resolution) was used to measure the cell geometry including the boundary and the out-of-plane cell height. IIT measurements were then performed on the same cell at multiple positions using multiple indentation depths. The FE model, created using the geometry from the LSCM measurements, was then used iteratively to determine the turgor pressure and wall modulus at each measurement position for which the computational data matched the measured data.

Fig. 2 provides an overview of the measurements. From the LSCM measurements, cell shape and the height of the pavement cell periclinal wall (for this example ∼3 μm) was measured relative to the average height of the surrounding anticlinal wall (Fig. 2A). The sample was then moved immediately to the IIT stage, and the same cell was identified (Fig. 2B). Several indentations (2.5 μm diameter tip) were made along the cell using the load function (Fig. 2C) developed previously (Forouzesh et al., 2013) that includes several short unloading segments at different depths. The shallow indentations were expected to be influenced mostly by the cell wall compression (Cosgrove, 2016) while deeper indents reflected a combination of the wall bending stiffness and turgor pressure. General IIT guidelines (Fischer-Cripps, 2011) suggest that indentation depths < 10 % of the sample thickness will be insensitive to any “substrate effect” which here refers to the turgor pressure. The minimum indentation depth is about half the thickness of the wall which means that the turgor pressure will have at least some effect on the measured response. The test positions were selected along the ridge of each cell near its maximum height for two primary reasons. First, measurements made along the cell plateau will have a reduced influence from the boundary conditions which are not known fully. Second, the ridge positions are primarily flat which reduces potential horizontal loads on the indenter tip to improve the reliability of the measurements. From each resulting force-displacement measurement (Fig. 2D) the local stiffness values (Fig. 2E) were determined. The stiffness values at all depths and the overall height of the periclinal wall were matched iteratively with the FE model of the experiment to find the turgor pressure and local wall modulus. For the FE simulation, the nanoindenter tip was measured and rendered within the FE software (additional details in Supplemental Figs. S1 and S2, Table S1). The measured cell boundary was used to create the cell model with wall thickness determined from high-resolution transmission electron microscopy (TEM) as shown in Fig. 3A-C. The surrounding anticlinal walls of the cell of interest were constrained on the outside by a stiff isotropic matrix to approximate the boundary conditions within the leaf. To understand the influence of surrounding cells, a set of expanded models was used as shown in Fig. 3D-F. The results provide a comparison between the IIT simulations for three cases: a single cell embedded in a matrix, a single layer of cells with neighboring cells, and a model with two layers of multiple cells. The influence of neighboring cells was most important for measurements near the boundary of the cell because the anticlinal wall was engaged during the deformation, but only for large indentation depths (> 2000 nm). However, for measurement positions near the maximum of the periclinal wall (central position Fig. 3D), the neighboring cells have a negligible effect. Therefore, the influence at the maximum depth used in the IIT experiments (≤ 1500 nm) was minimal (< 5 %). In addition, the deflection of the cell wall during the measurements will result in a decrease of the interior volume of the cell. The maximum percentage change in the volume was estimated to be less than around 0.5 % for the indentation depth of 1500 nm (Supplemental Fig. S5). These analyses were used to provide a reference to determine the experimental parameters of the approach in order to measure and evaluate the cell wall mechanical properties and turgor pressure accurately. Based on the experimental results, the simulation of the IIT measurements was used iteratively on the pressurized periclinal wall to estimate the mechanical properties at each position as well as the turgor pressure within the cell (results for all cells studied are shown in Supplemental Fig. S3; example comparisons between model and experiments are shown in Supplemental Fig. S4).

**Figure 2.**
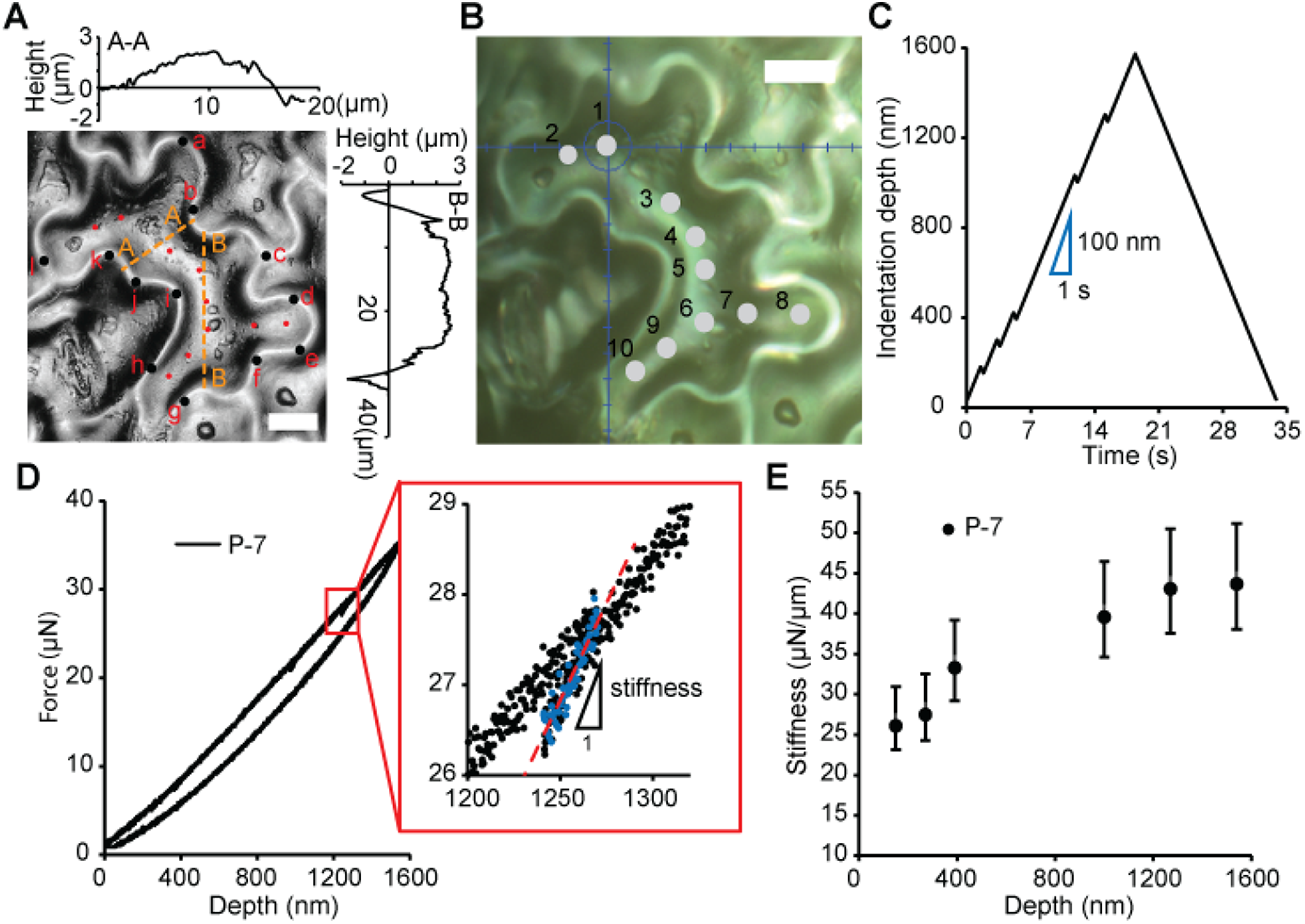
Geometric and mechanical characterization for turgid pavement cells in air. A, Topography of a pavement cell measured using a laser scanning confocal microscope (LSCM). The black points on the anticlinal wall were to calculate an average height as a reference, and the points on the periclinal wall (also shown in panel B) denote the positions of cell height measured relative to the reference. Two cross sections (A-A and B-B) show the measured height variation across this cell. B, Optical view of a pavement cell using the 50X objective of the indentation instrument. C, Input load function for the IIT experiments. The loading rate is 100 nm/s. D, Example measured force as a function of displacement and the associated stiffness. The stiffness is the slope of the unloading ramp and is used to determine the elastic modulus and turgor pressure. E, Stiffness at different depths at position 7 shown in panel B. Scale bar for (A) and (B) = 10 μm.

**Figure 3.**
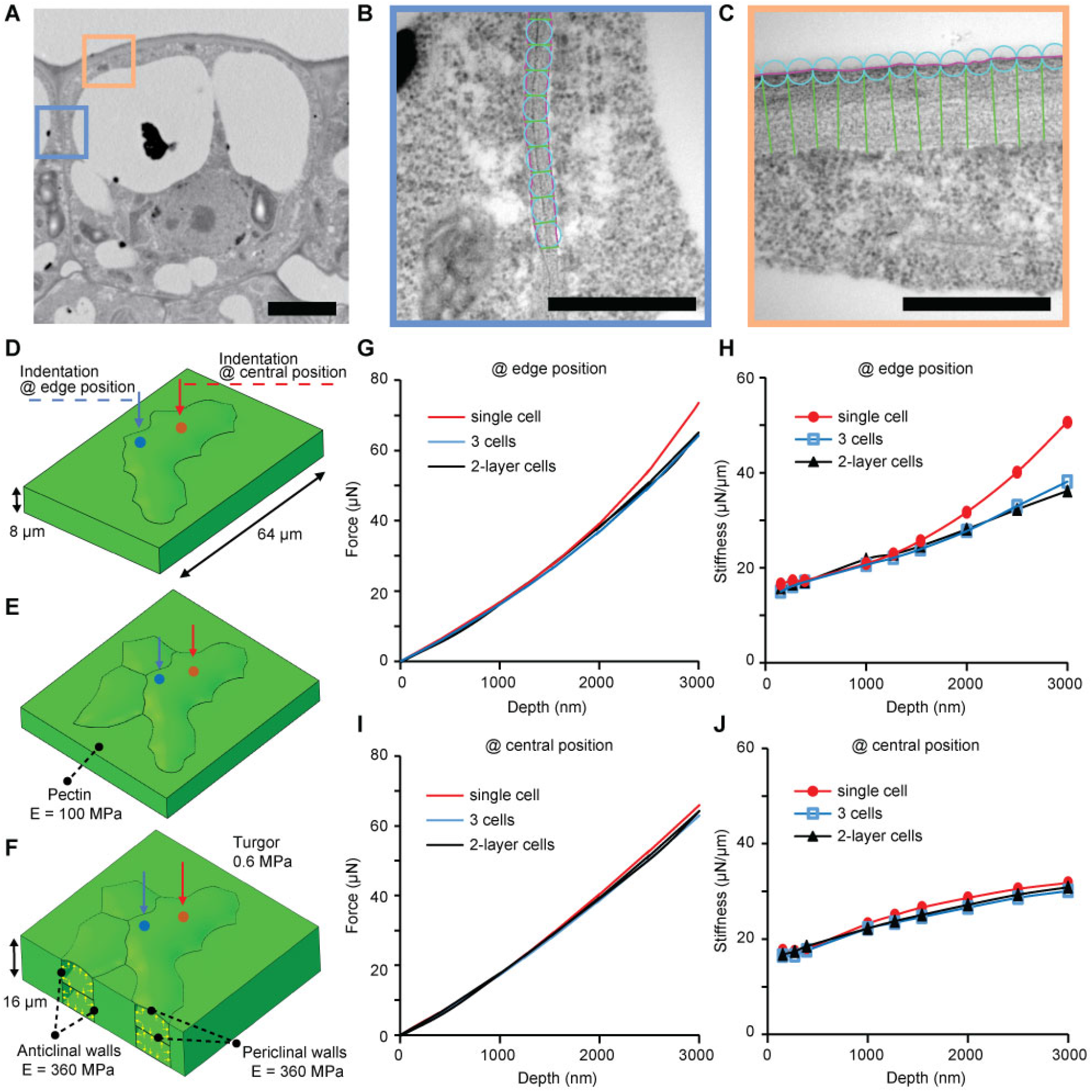
Details of geometry and the role of neighboring cells on the mechanical response from indentation. (A-C) TEM images show the cross-section of Arabidopsis cotyledon pavement cells (scale bar = 5 μm). The average thickness of the periclinal and anticlinal walls shown are 280 ± 14 nm and 35 ± 10 nm, respectively. (D) FE model of a single pavement cell. (E) Three cell structural model. (F) FE model with two layers of three cells. The cells at the bottom are assumed to be the same as the associated epidermal cells with an intermediate cell wall thickness of 50 nm. Simulations of IIT are performed, respectively, at the edge (4 μm away) and the central positions for all three models. (G, H) Force curve and estimated stiffness at different depths of the simulations indented at the edge position on a cell. (I, J) Simulation results for indentation at the central position of the cell. Note that the measurements described in the main text were limited to a maximum indentation depth of 1500 nm.

### Influence of cell geometry alone on the mechanical response

The IIT data (Fig. 4A) showed that the mechanical response at each indentation position along the periclinal wall was different. These differences reflect the combined influence of wall properties and cell geometry. To study the effect of cell geometry alone on the measured mechanical response, the FE model was used with a cell under an assumption of uniform mechanical properties across the periclinal wall (Fig. 4B). The mechanical response in the constricted region between two cell ingrowths was stiffer than that of other positions (Fig. 4C) due to the reduced distance between the sections of deformation. Additionally, the analysis showed that most material in the constricted regions had a similar mechanical response (P-2, P-3, and P-7 in Fig. 4C). Such an effect of geometry on the mechanical responses of pavement cells was expected. However, a comparison of the response with the IIT measurements, showed that cell geometry alone was not enough to explain the different responses for the five positions shown in Fig. 4A. For example, the measurement at position 7 was much stiffer than positions 2 and 3 which showed the same response in the model when uniform properties were assumed. Thus, the mechanical properties of the periclinal wall are clearly not uniform.

**Figure 4.**
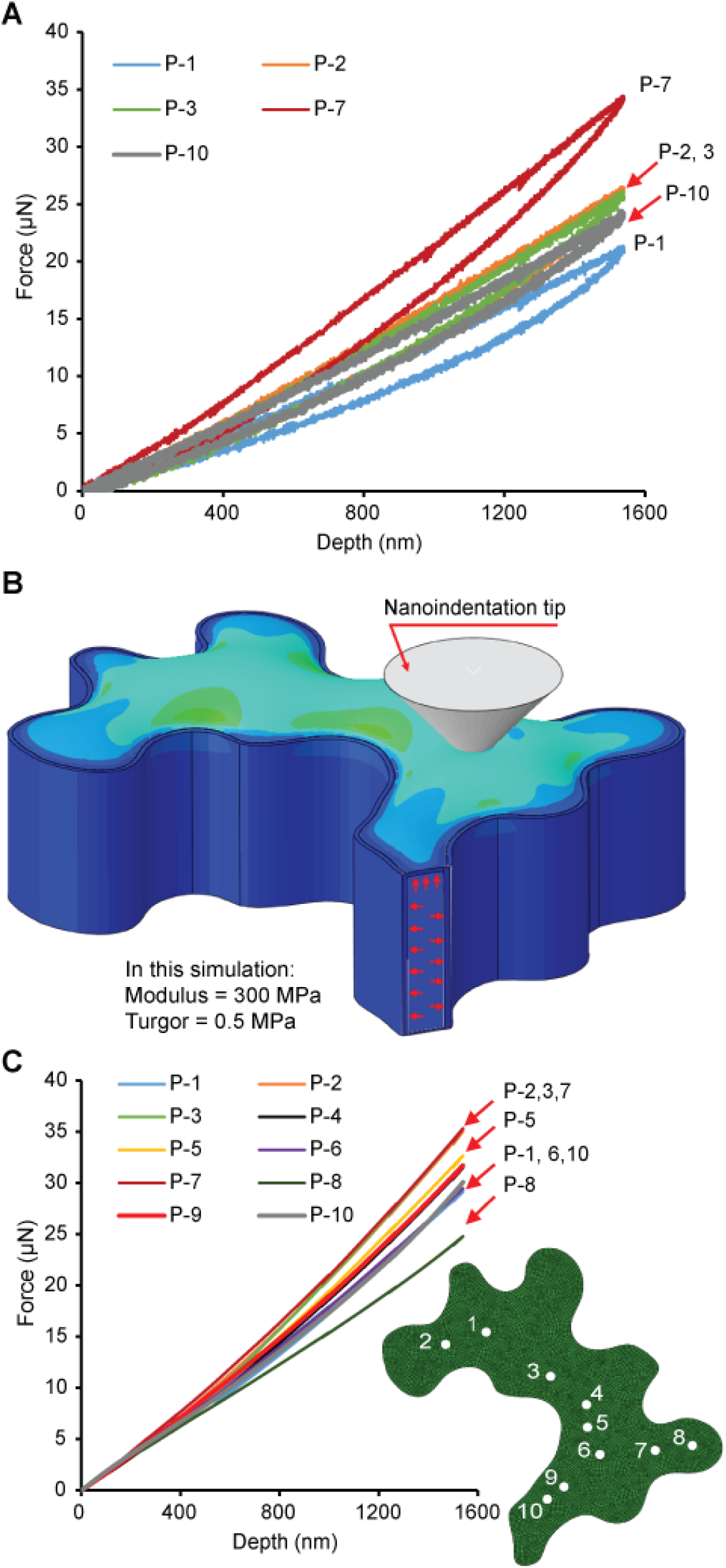
Influence of the cell geometry on the mechanical response. (A) Measured indentation force as a function of displacement at the ten positions of the periclinal wall shown in Fig. 1B. (B) Finite element model of the indentation experiments assuming uniform wall properties. The diameter of contact from the probe is ∼3 μm at the deepest position (see Fig. S5). (C) Indentation forces calculated using finite element analysis. A movie of the simulation is included in the Supplemental Material.

Using the iterative calculation based on the FE model (Supplemental Fig. S2), the local wall modulus and the associated turgor pressure at each indentation position of the periclinal wall were determined in order to match the geometric (i.e., out-of-plane periclinal wall deformation) and IIT mechanical measurements (see Fig. S4 for example comparisons between the final model result and the experimental data for the cell shown in Fig. 2). Results for four different cells are shown in Fig. 5. Deviations between the measurements and model results after iteration were very good and less than 20 % on average (see Table S6 for all results). The local elastic modulus across the wall shows a non-uniform distribution in the range of 200-800 MPa. Materials in several constricted regions were stiffer, but no clear trend was observed based on these measurements. For example, cell 1 (Fig. 5A-C) had moduli for constricted positions 1, 2, and 7 much higher than the rest (500-750 MPa), but other positions with constrictions had moduli closer to 400 MPa (positions 8, 9, and 10). For cell 2 (Fig. 5D-F), positions 4-6 had moduli greater than 450 MPa, while other positions were less than 400 MPa. For cell 4 (Fig. 5J-L), moduli in one constricted region (positions 4 and 5) were about half of the maximum > 600 MPa at position 2 which was more central. On the contrary, the turgor pressure estimated at each position in the cell was relatively uniform for each cell as expected. However, turgor pressure clearly varied for the different cells that were from different leaf samples, likely reflecting a variable water status for each leaf. Depending on the cell shape, size, and the growth conditions of the plants, the turgor pressure in most pavement cells at the status of the experiments here was expected to be on the order of 1 MPa (Forouzesh et al., 2013; Beauzamy et al., 2014) and the measurements agree.

**Figure 5.**
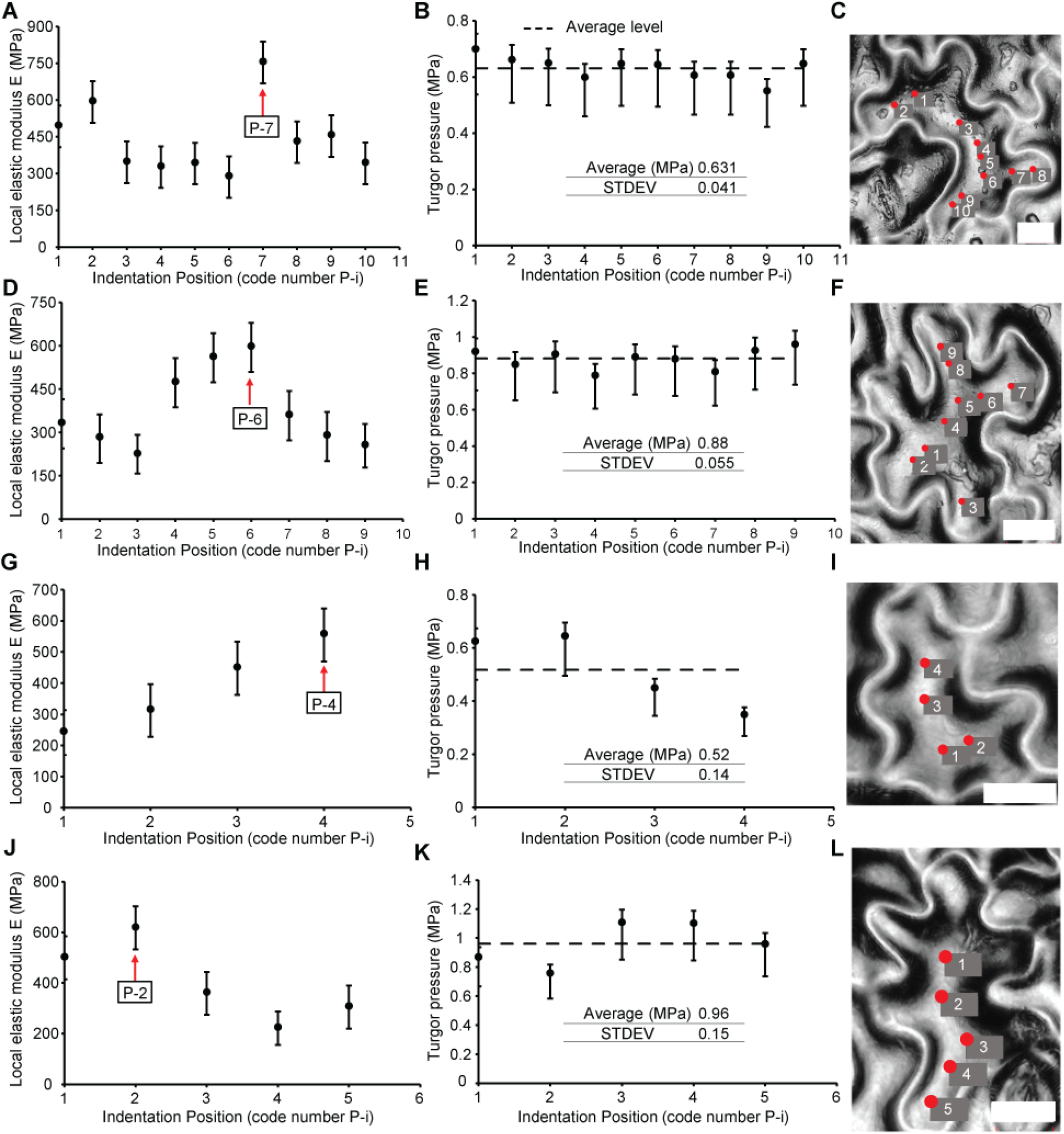
Wall modulus and turgor pressure results using the experimental-computational approach for four different cells. Elastic modulus (A, D, G, J) and turgor pressure (B, E, H, I) for each indentation position along the periclinal wall of the cells shown in the microscope images (scale bar = 10 μm). The horizontal axis represents the indentation position (defined in the microscope images). The uncertainty in the plots is estimated by considering the uncertainties in the measured height and stiffness data as shown in Fig. 2A and E.

### Mechanical response for different osmotic conditions

The turgor pressure values were validated using measurements on a cell subjected to different osmotic conditions. Mannitol solution treatments (Fig. 6A; Methods) were used to manipulate the turgor pressure in cells (Lucas and Alexander, 1981). As the concentration of the solution increased up to 0.9 MPa solute potential, the relative height of the periclinal wall was relatively constant with a slight trend downward. The height decreased dramatically after a solute potential of 1.2 MPa (Fig. 6B and C). Optical images of the cell at different stages are shown in Supplemental Fig. S6. As expected, the IIT experiments showed the effect of reduced turgor pressure. The mechanical force in the cell wall was reduced (Fig. 6E), and for the range of 0.9-1.2 MPa solute potential, the mechanical response no longer changed. The highest concentration solution (1.5 MPa) greatly modified the surface quality of the cell, such that IIT experiments could no longer be conducted due to the difficulty of cell identification.

**Figure 6.**
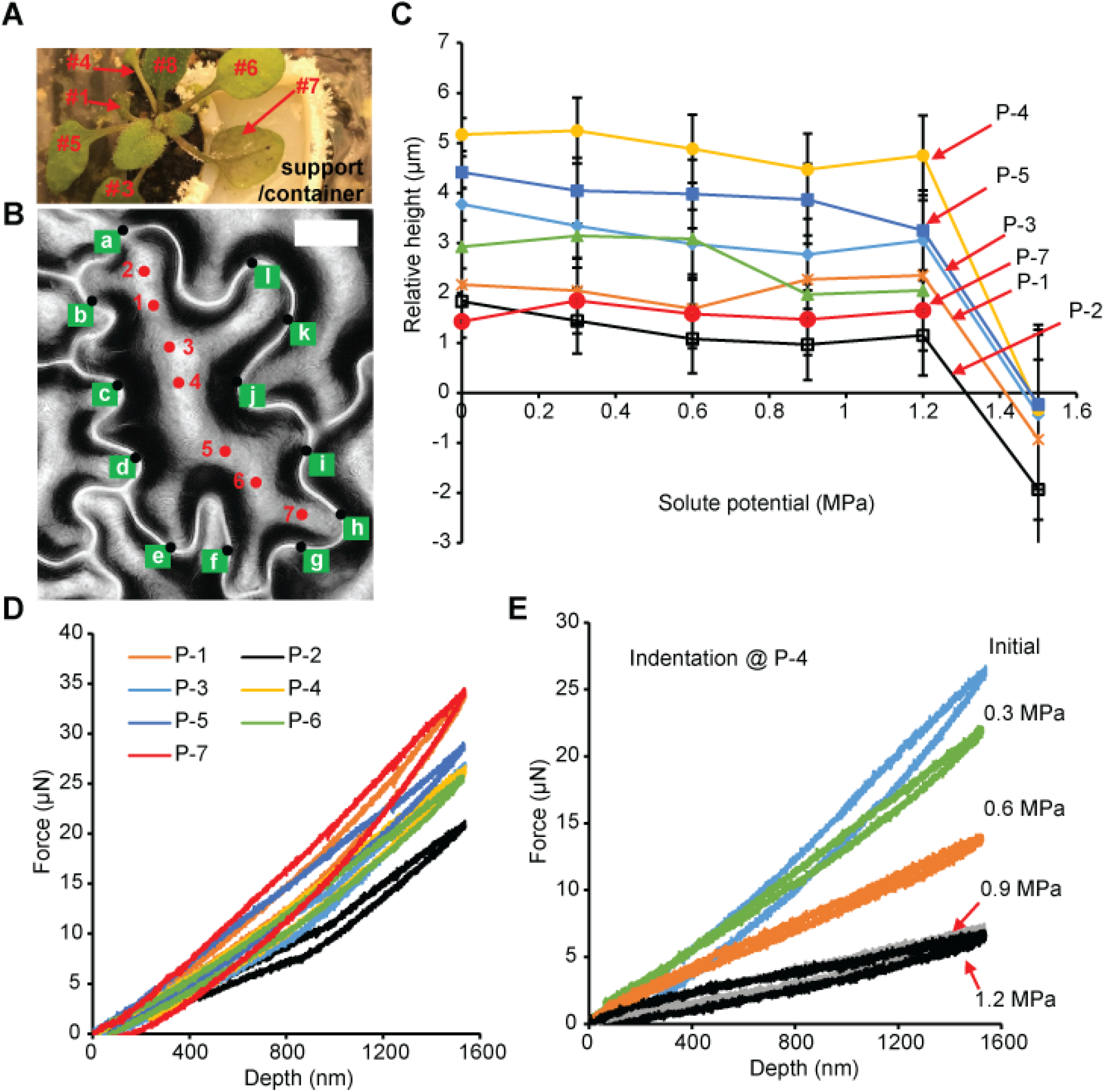
Geometric and mechanical characterization of the periclinal cell wall for different osmotic conditions. (A) Experiment setup. The adaxial side of a leaf was glued on a support and immersed in different concentrations of D-mannitol solution. (B) Height measurement of the periclinal wall. The green points on the anticlinal wall are used for calculating the average boundary height as a reference, and the red points are the positions of cell height measured relative to this reference. Scale bar = 10 μm. (C) Relative height of the periclinal wall as a function of solute potential. (D) Indentation force vs. displacement across the periclinal wall at the initial stage before mannitol treatment. (E) Indentation force vs. displacement for position P-4 for different mannitol treatments.

At each stage of the solution treatment, IIT measurements were made only at positions P-3, P-4, and P-5 in order to reduce the overall measurement time for each stage of treatment. As a result, estimated changes in turgor pressure and modulus were made only at these positions (all measurement data are shown in Supplemental Fig. S7). Fig. 7A shows that turgor pressure in the pavement cell dropped with increasing of concentration of the mannitol solution. The turgor pressure for this cell at the initial stage was estimated to be 0.8 MPa, and the reduction in turgor pressure agreed well with the difference of solute potential between the first two treatments. For the remaining treatments, the pressure decreased nonlinearly until dropping to the range of 0.1-0.2 MPa at the associated conditions of 0.9 and 1.2 MPa solute potential. This minimum value of the turgor pressure from the IIT measurements was on the order of the measured variation in the turgid cells. The distribution of elastic modulus along the periclinal cell wall (Fig. 7B) was non-uniform and was similar to the results from other cells (Fig. 5). However, the local elastic modulus was relatively constant (< 15 % drop) at indentation positions P-3, P-4 and P-5 as the turgor pressure changed (Fig. 7B). For higher solution concentrations, the elastic modulus of the cell wall appeared to degrade slightly although these data are insufficient to make any strong conclusions.

**Figure 7.**
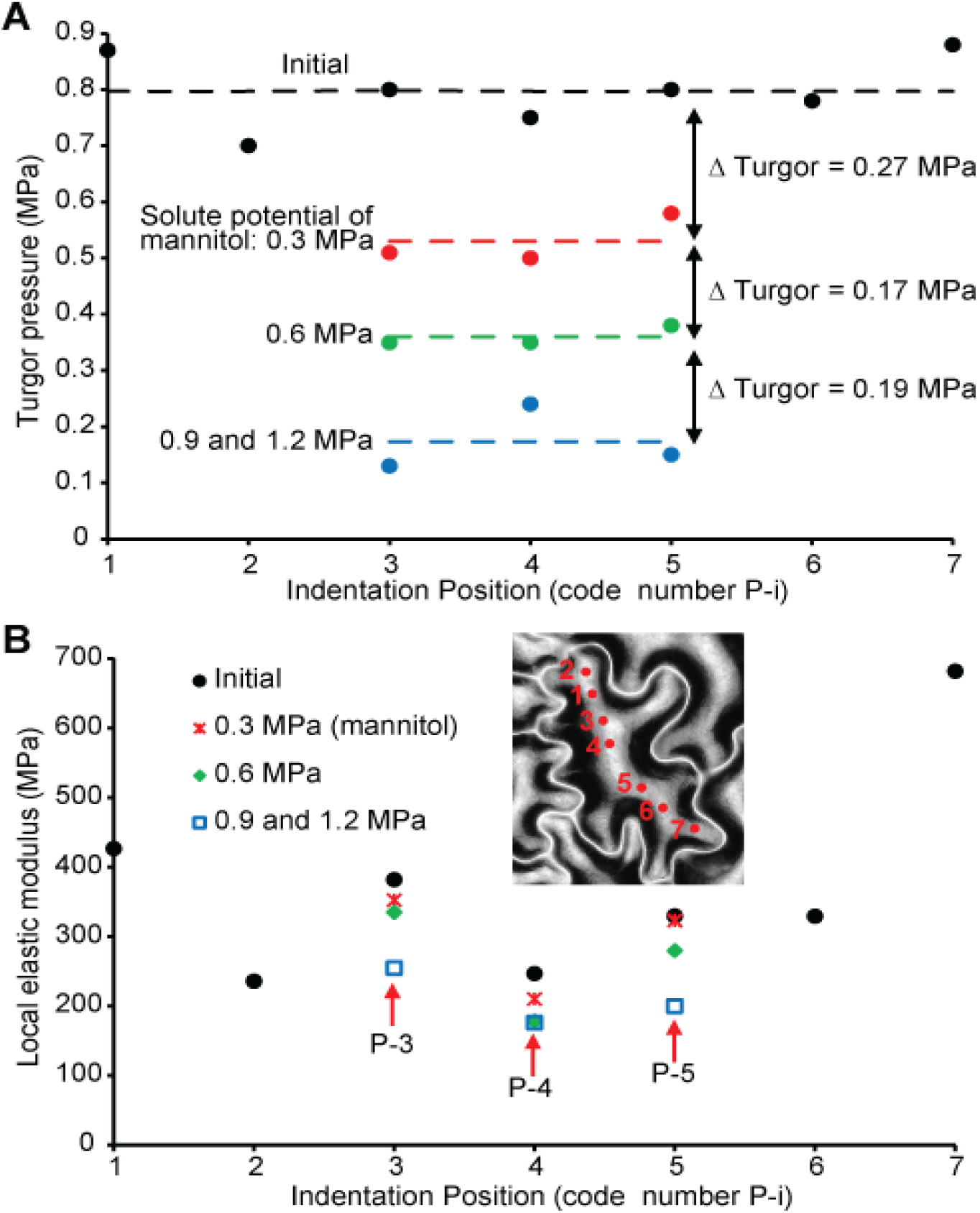
Results using the iteration method for the pavement cell under different osmotic conditions. (A) Turgor pressure in the cell estimated at each indentation position for different mannitol solution treatments. (B) Elastic modulus of the cell wall at each indentation position as a function of solute potential treatment.

## Discussion

The integrated experimental-computational approach described here used both optical (LSCM) and IIT measurements. This combination allowed both the deformed shape of the wall and several force-displacement measurements to be used in the model. The iterative approach provided local values of wall modulus and turgor pressure. The results showed that the mechanical response varied at different positions across the cell although the turgor pressure values were very uniform. The FE simulations of the measurements with uniform cell wall properties showed that cell geometry alone cannot explain the measured variations. The cells studies were relatively mature (22-28 days after germination) such that they were likely fully lobed. Thus, the variation in modulus must have occurred during the growth process which is counter to simplistic models for cell lobing control based solely on geometry (Sapala et al., 2018). Previous results (Sampathkumar et al., 2014) reported enhanced stiffness in the convex region of lobes, but their AFM protocol did not deform the periclinal wall to the extent used here. Our results show that the convex regions were not always stiffer than other positions. The maximum load of the measurements was 40 μN, 40 times the maximum used in Sampathkumar et al., 2014, such that the mechanical deformation engaged the full thickness of the periclinal wall in bending (a movie of a simulation is included in the Supplemental Material).

Pavement cell expansion during growth is known to occur non-uniformly based on measurements of cell wall deformation using time lapsed quantification of cell shape (Zhang et al., 2011; Wu et al., 2016) and microbeads to measure local growth behavior (Armour et al., 2015; Elsner et al., 2018). Different cells also grow at different rates and a single cell may have walls that each expand at different rates (Belteton et al., 2021). Cell enlargement is related to the change in strain of the periclinal wall and anticlinal walls, and non-uniform growth must have a non-uniform strain distribution across the cell walls. Sapala et al., 2018 proposed that geometry alone is sufficient to predict the growth patterns of pavement cells but our results suggest that material property variations are also important. A clear understanding of the distribution of material properties within single cells is needed for robust growth models. The properties may change over time and the approach described here can be used to quantify and track such changes.

The range of the elastic modulus of the periclinal wall of *Arabidopsis* pavement cells varies from 200-800 MPa, and this range is consistent with previous results based on IIT (Forouzesh et al., 2013). The sensitivity study for the modulus estimate based on a uniform thickness (Supplemental Table S2) shows that variations of cell wall thickness could play a role in the outcome. The TEM measurements showed a small deviation of ∼20 nm (Fig. 3C), but variations on the order of 50 nm could affect the modulus estimates by ∼30 %. The variation of modulus within a single cell could reflect a variation in volumetric content (i.e., the percentage of cellulose at a given position), but is more likely a reflection of local anisotropy from cellulose microfibril alignment. The FE model used here is based on isotropic material behavior so these results will direct future models that can incorporate in-plane anisotropy. The modulus values here are much larger than some literature reports, primarily those based on AFM measurements (Peaucelle, 2014; Sampathkumar et al., 2014; Beauzamy et al., 2015). In some cases, moduli less than 1 MPa have been used in FE models (Sampathkumar et al., 2014; Majda et al., 2017; Bidhendi et al., 2019). Modulus values in this range would lead to significant expansion of the periclinal wall (as shown in the example FE result of Fig. 1E) unless walls are thicker or the values of turgor pressure are much lower. An AFM tip would engage only a small volume of material at the outer surface of the wall (Cosgrove, 2016), which means that such results are not expected to match IIT measurements that bend the entire cell wall. Moduli of several hundred MPa are consistent with a composite that includes both cellulose, with stiffness of tens of GPa, and a viscoelastic matrix comprised of pectin, hemicelluloses, and numerous proteins.

The turgor pressure values were found to be nearly constant for all measurements within a given cell. This outcome is expected (Beauzamy et al., 2014). However, there was variation between cells from different leaves, even though the age of all leaves was similar. These differences likely reflect the varying water status of the leaf. However, very little information is found in the literature about turgor pressure variations across a single leaf primarily because such measurements have historically been destructive and have not been possible on small cells. Here, turgor pressure measurements after the first two stages of mannitol solution treatment showed a decrease that was consistent with the difference of the two solute potentials. Such results highlight the sensitivity of the measurements to small changes in turgor pressure. The changes in turgor pressure for the higher solute potentials reflect the nonlinear behavior of the cell volume change that has been observed in connection with solution concentration (Sajnin et al., 1999). These results validate the turgor pressure estimates using the IIT-based approach.

Using higher concentrations of mannitol, turgor pressure in cells decreases, whereby the periclinal wall gradually flattens. For osmotic potentials of 0.9-1.2 MPa, the height of the cell wall shows a small change, but is followed by a dramatic drop. These stages are associated with the presence of plasmolysis. This phenomenon is also consistent with previous reports (Speth et al., 2009). Generally, the plasmolysis starts in some locations of the cell for mannitol solutions of 0.6-0.8 mM (1.48-1.98 MPa solute potential) (Lang et al., 2014; Junková et al., 2018). The extent of the changes during mannitol treatment is comparable with the threshold value of solute potential for plasmolysis which was in the range of 1-2 MPa (see Supplemental Fig. S8). This information supports the estimation of turgor pressure using our approach.

Our measurements also show that the elastic modulus remained constant for the first two stages of mannitol treatment, but decreased by ∼35% for higher mannitol concentrations. A similar trend was reported by (Beauzamy et al., 2015), but they found a decrease of modulus of ∼85% for high concentrations of sorbitol solution which they attributed to a drop in wall tension from the pressure. Our model accounts for such wall tensioning which may explain the smaller reduction of modulus. Another previous result (Forouzesh et al., 2013), in which a salt solution was used to reduce turgor pressure, did not show significant differences in wall modulus due to plasmolysis, but their lack of sensitivity may have been the result of limited information of cell geometry used in their model. Some studies have shown that a saline solution can lead to an increase in transport of Na^+^, K^+^ and Ca^2+^ out of the cell (Cramer et al., 1985; Almeida et al., 2017). It has also been shown (Zsivanovits et al., 2004) that the loss of these elements can lead to a stiffness reduction in pectin. Thus, the water movement may also cause ion transport out of the wall matrix leading to changes that are detectable for higher concentrations of the mannitol solution. However, cellulose is the primary load-bearing component of the wall so small changes in natrix modulus may not be measurable for small values of wall strain. In general, the use of chemical treatments to manipulate turgor in cells remains a valuable technique but more studies are needed to assess potential changes to cell wall mechanical properties.

## Conclusions

The integrated experimental-computational approach described here allows accurate 3D geometry of the cell to be used to create a robust FE model to simulate the indentation experiments that deform the ***entire thickness*** of the cell wall. The approach has sufficient spatial resolution to map the properties of the periclinal wall on living *Arabidopsis* pavement cells. The results quantitatively demonstrated a variation of properties within single cells and is sensitive to wall material throughout the entire thickness, an aspect of this work which is important for future cell growth and development studies. Even though anisotropic mechanical properties are not considered in this article, these results serve as a reference for additional studies to characterize the in-plane anisotropy ratio in the periclinal wall, in order to understand the morphogenesis of the epidermal cells during growth (Belteton et al., 2021). This method was also used successfully to estimate turgor pressure for living plant pavement cells nondestructively. It has the potential to be used to monitor the change of turgor for the pavement cells during growth and under specific chemical or environmental conditions.

## Materials and methods

### Plant materials

*Arabidopsis thaliana* wild type (*Col-0*) was used for the measurements. The plants were grown in a growth chamber (23°C, 16 hours light/8 hours dark), and 22∼28 days after germination, the plant was transplanted in a small petri-dish attached with a support and settled for 1∼3 days before testing. Leaves (#5, #6, or #7) were mounted on the support using epoxy gel (5 min cure, Devcon) to expose the adaxial side. The soil in the petri-dish was covered with plastic film to prevent dehydration. To be able to identify the same epidermal cells, a black Sharpie marker was used to make a reference mark on the sample surface (a very small mark away from the test site). Five plants were used for the experiments involving a single leaf from each plant. Trichomes sometimes limited the cells that could be studied, but were not problematic.

The Arabidopsis line expressing TUB6:GFP having green fluorescent protein (GFP) tag was used to observe plasmolysis under different osmotic conditions. The #5-#7 leaf (30-40 days after germination) of the plant was cut, and the petiole was sealed using epoxy. Then the adaxial side was flattened and glued on a thin glass slide for microscopy.

### Mannitol solution preparation

D-Mannitol was purchased from Sigma Aldrich (type number M4125), and different solute concentrations of the solution (D-Mannitol and distilled water), 0.12 M (0.3 MPa), 0.24 M (0.6 MPa), 0.36 M (0.9 MPa), 0.48 M (1.2 MPa) and 0.61 M (1.5 MPa), were prepared. The numbers in parentheses are the associated solute potentials (*ψ*_*s*_) which are calculated using*ψ*_*s*_ = −*icRT*, where *i* = 1 for mannitol, *c* is the solute concentration (unit is mol/L), *R* is the pressure constant (8.314 kPa**·**L/mol**·**K) and *T* is the absolute temperature.

### Mannitol solution treatment of the samples

Two groups of samples were prepared for mannitol treatment. For the first group, there was no treatment, and the living leaf epidermis was mounted directly on a support for optical and mechanical testing. For the second group, a plant leaf was mounted on a support/container, and each mannitol solution was placed in the support/container for 50 minutes to allow the water potential in the epidermal cells to reach equilibrium. Then the solution was carefully removed from the container, the leaf was carefully blotted dry, and the LSCM and IIT measurements performed.

### Laser scanning confocal microscope and TEM experiments

A laser scanning confocal microscope (LSCM) (Keyence VK-X200, 402 nm wavelength, 0.5 nm z-resolution) with an objective lens of 50X (long working distance) was used for measuring the 3D shape of the epidermal cells tested in this study. Using this microscope, the boundary geometry of the cells of interest and the relative height of the periclinal wall relative to the average height of the surrounding anticlinal wall were measured.

Another confocal microscope (Zeiss LSM 800 with Airyscan, Germany) was used to observe plasmolysis of the GFP samples. The green channel and the 50X and 10X objective lenses were used. For application of the 50X objective, oil was added on the lens for the optical observation of the same cells. The transmission electron microscopy (TEM) methodology described previously (Yanagisawa et al., 2015) was used to determine the thickness of the periclinal wall used in the FE model.

### IIT experiments

Instrumented indentation testing (IIT) experiments (Fischer-Cripps, 2011) were conducted in quasi-static mode using a Hysitron Triboscan Ti950 (USA) with a cono-spherical indenter tip (2-3 μm maximum contact diameter). A 50X magnification objective lens (long working distance) was used to observe the epidermal cells. All IIT measurements were conducted in air. Before testing, the leaf sample was settled for 1∼2 hours in the machine for equilibration. A force of 2-5 μN was used to engage the sample. Displacement control was used for the experiments. Before the experiment of each plant pavement cell, a tip-to-optic calibration (Fischer-Cripps, 2011) was conducted in order to ensure that the spatial positioning on each cell was as accurate as possible.

### Finite element method (FEM) and iterative calculation

The structural model of the epidermal cells was constructed, and the mechanical analysis was performed using commercial finite element software Abaqus (2019 version). In the FE models, the thickness of the periclinal wall was fixed at 300 nm based on TEM images (Fig. 3C), and the cell wall was meshed using elements (element type C3D8R) with a maximum size of 100 nm to ensure a minimum effect from the meshing. Each single periclinal wall was tied with the surrounding anticlinal wall, and the anticlinal wall was confined as a boundary condition. Except for the models with multiple cells (Fig. 3E and F) whose anticlinal wall thickness was 50 nm, the anticlinal wall thickness was fixed at 500 nm which played a minor role due to the surrounding matrix that was used to confine wall anticlinal wall deformation. Before pressurization, the periclinal wall was assumed flat. The material of the cell wall was assumed to be an isotropic neo-Hookean material, and the whole material was assumed to be a standard linear solid with a primary relaxation time of 6.8 s (Zsivanovits et al., 2004; Forouzesh et al., 2013). The influence of the relaxation time on the estimations of modulus and turgor pressure was very minor (< 5.3 %) for changes covering 4 times (supplemental Table S3-4). The role of relaxation/creep was larger (maximum ∼30 %) when the ratio *G*_i_/*G*_0_ was varied by a factor of 10 (supplemental Table S5). The Poisson’s ratio was assumed to be 0.47 (i.e., nearly incompressible). The equations for the mechanical model are described in the Supplementary Material. The cell wall was first pressurized and then the indentation simulation was performed.

For estimating turgor pressure and local elastic modulus at each indentation position, the hyperelastic material was assigned uniformly across the whole cell wall. Similar to the previous approach (Forouzesh et al., 2013), the stiffness at the shallow indentation depth is dominated by the elastic modulus but the turgor pressure affects the stiffness more at deeper measurements. Therefore, the simulated stiffnesses at all indentation depths were used to match those of the IIT experiments. Furthermore, at the end of cell pressurization, the vertical displacement of the wall at the indentation position was used to compare with the optical height measurement. Once both the optical and mechanical measurements are matched iteratively (refer to Supplemental Fig. S4), the turgor pressure and the local wall modulus were determined. The flowchart of the iterative approach is shown in Supplemental Fig. S2.

## Supplementary Data

Description of the mechanical modeling and parameters used in the FE analysis. Supplemental Figure S1. Standard linear solid model.

Supplemental Table S1. Parameters used in the FEM.

Supplemental Table S2-S5. Sensitivity study for the estimated turgor pressure and elastic modulus using, respectively, fixed thickness, relaxation time and Gi/G0 ratios.

Supplemental Figure S2. Flowchart for iteratively estimating turgor pressure and local elastic modulus in the periclinal wall.

Supplemental Figure S3. Nanoindentation experiments and height topography measurements of the pavement cells.

Supplemental Figure S4. Comparison of the simulation with the experiments.

Supplemental Table S6. Percentage difference between the experimental values and the final model values after iteration for all the cells.

Supplemental Figure S5. Change in the total cell volume for different indentation depths as calculated from the indentation model.

Supplemental Figure S6. Laser scanning confocal microscope images of the pavement cell at the stage of each mannitol solution treatment.

Supplemental Figure S7. Nanoindentation experiments on the pavement cell for different osmotic conditions.

Supplemental Figure S8. Optical observation of plasmolysis of the GFP leaf sample.

## Acknowledgements

This work was supported by the US National Science Foundation under grant number MCB-1715444 to D.B.S and J.A.T. The IIT experiments and LSCM measurements were conducted at the Nano-Engineering Research Core Facility (NERCF) of the University of Nebraska-Lincoln, which is partially funded by the Nebraska Research Initiative.

